# A comparison of super-resolution microscopy techniques for imaging tightly packed microcolonies of an obligate intracellular bacterium

**DOI:** 10.1101/2024.08.12.607698

**Authors:** Alison J. North, Ved P. Sharma, Christina Pyrgaki, John Lim S.Y., Sharanjeet Atwal, Kittirat Saharat, Graham D. Wright, Jeanne Salje

**Affiliations:** Bio-Imaging Resource Center, The Rockefeller University, New York, NY, USA; A*STAR Skin Research Labs (A*SRL), Agency for Science, Technology & Research (A*STAR), Singapore; Lonza, 8830 Biggs Ford Road, Walkersville, MD 21793, USA; Cambridge Institute for Medical Research, University of Cambridge, UK; Research Support Centre (RSC) Agency for Science, Technology & Research (A*STAR), Singapore; Mahidol-Oxford Tropical Medicine Research Unit, Faculty of Tropical Medicine, Mahidol University, Bangkok Thailand; Department of Biochemistry, University of Cambridge, UK; Department of Pathology, University of Cambridge, UK

**Keywords:** Super-resolution imaging, *Orientia tsutsugamushi*, Rickettsiales, obligate intracellular bacteria, STED, iSIM, 3D-SIM, confocal, Airyscan, Deep learning, 3D cell segmentation

## Abstract

Conventional optical microscopy imaging of obligate intracellular bacteria is hampered by the small size of bacterial cells, tight clustering exhibited by some bacterial species and challenges relating to labelling such as background from host cells, a lack of validated reagents, and a lack of tools for genetic manipulation. In this study we imaged intracellular bacteria from the species *Orientia tsutsugamushi* (Ot) using five different fluorescence microscopy techniques: standard confocal, Airyscan confocal, instant Structured Illumination Microscopy (iSIM), three-dimensional Structured Illumination Microscopy (3D-SIM) and Stimulated Emission Depletion Microscopy (STED). We compared the ability of each to resolve bacterial cells in intracellular clumps in the lateral (xy) axis, using full width half maximum (FWHM) measurements of a labelled outer membrane protein (ScaA) and the ability to detect small, outer membrane vesicles external to the cells. We next compared the ability of each technique to sufficiently resolve bacteria in the axial (z) direction and found 3D-STED to be the most successful method for this. We then combined this approach with a custom 3D cell segmentation and analysis pipeline using the open-source, deep learning software, Cellpose to segment the cells and subsequently the commercial software Imaris to analyze their 3D shape and size. Using this combination, we demonstrated differences in bacterial shape, but not their size, when grown in different mammalian cell lines. Overall, we compare the advantages and disadvantages of different super-resolution microscopy techniques for imaging this cytoplasmic obligate intracellular bacterium based on the specific research question being addressed.

## INTRODUCTION

Over the past 25 years, light microscopy of model bacterial organisms including *Escherichia coli, Bacillus subtilis* and *Caulobacter crescentus* has revolutionised the understanding of bacterial cells, revealing them as highly organised, with a cytoskeleton and dynamic molecular machines involved in DNA segregation and cell division^1,2^. This new paradigm has been based on conventional widefield and confocal microscopy^3,4^ as well as newer super-resolution techniques^2,5^, combined with labelling approaches including immunofluorescence, genetically encoded fluorescent proteins, and small fluorescent tags. Almost all the resulting knowledge on bacterial growth and division is derived from free living bacteria, because of challenges with applying these techniques to intracellular bacteria, especially obligate intracellular bacteria that cannot be manipulated axenically. Specific challenges include the genetic intractability of many obligate intracellular bacteria, their reduced cell size compared with free living counterparts, and their tendency to grow in tightly-packed three dimensional bacterial microcolonies^6–9^.

Orientia tsutsugamushi (Ot), an obligate intracellular bacterium belonging to the Order Rickettsiales and Family Rickettsiaceae, presents these challenges when attempting to accurately determine cell morphology using conventional imaging methods. Approaches to date have primarily involved immunofluorescent labelling of the major surface protein TSA56, followed by widefield or confocal microscopy^6–8^, although the use of the super-resolution techniques of STORM and SIM imaging has been reported^9,10^. Ot exhibits pleomorphism, with individual cells adopting rod, round, or irregular shapes, as documented through confocal and electron microscopy^8,9^. However, comprehensive and systematic studies regarding the distribution of bacterial morphologies in this species are lacking due to the inherent difficulty in precisely measuring the cells in three-dimensions. Bacterial shape is primarily governed by a rigid peptidoglycan cell wall, shaped by an intricate internal cytoskeleton. This three-dimensional architecture significantly impacts bacterial fitness in various environments, influencing factors such as nutrient uptake efficiency, motility patterns, and susceptibility to predation by phagocytes or bacteriophages. Our study aims to compare the ability of a range of super-resolution microscopy techniques to resolve individual Ot in intracellular clumps to enable the accurate quantification of the three-dimensional morphology of Ot within infected host cells. Such an investigation holds promise for advancing our understanding of the role of bacterial shape in Ot fitness and pathogenicity.

Ot causes the mite-borne human disease scrub typhus that is endemic in Asia, with closely related species recently described in other parts of the world including Latin America and the Middle East^11^. There is currently no vaccine available and the mean mortality rate in untreated cases is 6% making it one of the most severe rickettsial infections in humans^12^. Ot can replicate in a range of cell types including endothelial, fibroblast, epithelial, monocyte/macrophage and dendritic cells^8,13,14^. Ot enters cells using clathrin-mediated endocytosis^15^ and macropinocytosis^9^, then traffics to the perinuclear region where it undergoes replication to form a tightly packed bacterial microcolony^6,9^. Effective imaging, segmentation and analysis of individual Ot bacteria in this perinuclear microcolony is limited when using traditional labelling and imaging techniques.

Many light microscopy techniques capable of imaging specimens at resolutions exceeding the classical Abbe diffraction limit have emerged over the past two decades (see a variety of excellent papers including ref.^16–20^). These super-resolution methods include Stimulated Emission Depletion Microscopy (STED^18,21–23^), single molecule localisation techniques like Stochastic Optical Reconstruction Microscopy (STORM^24^), and a range of Structured Illumination Microscopy (SIM)-based techniques^17,25^ that encompass both conventional 3D-SIM^16,26,27^ and more recent image scanning microscopy approaches^28^ including instant SIM (iSIM)^29^. In addition, techniques to enhance the resolution of confocal microscopy have become widely used^17,30^, either through combining highly sensitive detectors with a reduced confocal pinhole diameter, or by the integration of specific detectors that use pixel reassignment methods to increase sensitivity and resolution, such as the Zeiss Airyscan detector^31^ and, more recently, the Abberior Matrix detector^32^. While super-resolution methods initially relied upon the use of home-built and typically user-unfriendly systems, the availability of excellent commercial products, especially when housed within dedicated microscopy core facilities supported and operated by expert staff experienced in their application and optimization for varied samples, has rendered these techniques more widely accessible to research biologists^33^.

The goal of this study was to compare the ability of different super-resolution microscopy techniques to image intracellular Ot and develop custom analysis pipelines for 3D bacterial cell segmentation and quantification of cellular parameters. We compare four different super-resolution microscopy techniques with each other and with conventional confocal microscopy (given the much wider familiarity with this technique) and present the advantages and disadvantages of the respective techniques for studying Ot. Importantly, all selected techniques were applied using commercially available instruments, which were in general operated using the manufacturer’s recommended acquisition settings, to enable researchers elsewhere with equally challenging biological samples to apply a similar approach.

The microscope techniques compared here were: standard confocal microscopy using a Zeiss LSM 880 system (“confocal”); Airyscan confocal microscopy using a Zeiss LSM 880 system fitted with an Airyscan 1 detector (“Airyscan”); 3D-Structured Illumination Microscopy using an OMX V4 Blaze microscope (“3D-SIM”); instantSIM using a VisiTech Vt-iSIM system (“iSIM”); and Stimulated Emission Depletion Microscopy using an Abberior Facility Line system (“STED”). The exact specification of each microscope system is detailed in the Methods section. It should be noted that the point spread function (PSF) of the Abberior Facility Line STED system used here can be shaped, and optimized, in both the lateral (xy) and axial (z) axes^18,34^, thus when we use the term “3D-STED”, we are referring to a PSF shaped in all 3 dimensions, not simply to the acquisition of a z-stack of images acquired with a 2D-shaped PSF. The percentage of 2D-STED vs. 3D-STED can thus be easily tuned by means of a simple slider either to maximize the xy resolution (using 100% 2D-STED) or to give near isotropic resolution (100% 3D-STED), or a compromise between the two, an option that is unique to STED amongst super-resolution methods. Thus, various ratios of 2D:3D STED were tested here. Sample labelling and choice of fluorochrome was optimized for each technique. Whilst labelling precision could be further enhanced by using a method with lower linkage errors (nanobodies, directly labelled FAb fragments, self-labelling protein tags, etc.), indirect anybody labelling was a simple approach and quite adequate for the level of resolution we needed to achieve^35^.

## RESULTS

### Multiple imaging platforms enabled bacteria to be resolved in two dimensions, but only 3D-STED imaging could fully resolve them in the axial direction

We compared the ability of five imaging techniques to image and resolve fixed, immunolabelled Ot bacteria within intracellular microcolonies. Secondary antibodies were selected using dyes previously shown to be optimal for each microscope technique. For classic fluorescence techniques such as confocal, and enhanced resolution or super-resolution techniques whose resolution is still fundamentally bound by the laws of diffraction^34^, including 3D-SIM, Airyscan and iSIM, the resolution that can be achieved depends directly on the wavelength of fluorescence emission, with resolution being higher with shorter wavelengths (see Jonkman et al.,^36^ for a practical example). The actual resolution achieved with methods such as 3D-SIM is also strongly dependent on the dye’s brightness and resistance to photobleaching, since these factors directly affect the quality of the reconstruction, as well as by the acquisition and implementation of high quality, empirically determined PSFs^37^. Thus, we tested samples labelled both using AlexaFluor (AF) 488-conjugated secondaries and DyLight 405-labelled secondaries, expecting that the 405-labelled samples should give the highest resolution. AF488 is a very commonly used dye, with a deservedly strong reputation for being both bright and highly photostable. In contrast, few investigators use blue-emitting dyes because of their reputation for being weak and easily photobleached and given the challenges of natural tissue autofluorescence in this region of the spectrum. However, previous experiments comparing the combination of different commercial blue-emitting dyes with a variety of antifade-mountants have demonstrated that DyLight 405-conjugated antibodies, combined with Prolong Diamond mounting medium, are sufficiently bright and photostable to achieve high quality images of many cellular structures with 3D-SIM (A. North, unpublished data).

For STED microscopy, resolution is dependent on the photophysics of the fluorochrome rather than its wavelength - it must be bright, photostable, and easily driven down into a non-emitting state by the depletion laser (reviewed by Gould *et al*.^38^). In others’ work and also our hands, both the far-red dye STAR-RED, developed by Abberior specifically as a STED dye^39–43^, and the more conventional AlexaFluor dye AF594 have given excellent results in STED microscopy^44–46^, working well in combination with the 775 nm depletion laser on the Abberior Facility Line microscope. Thus, we selected these two dyes for comparison here. Both dyes enabled us to resolve the bacteria using 3D-STED imaging, but the results obtained with STAR-RED were superior, hence we only feature STAR-RED labelled samples here and used this dye alone for the remainder of the STED imaging.

Figure 1 shows examples of the best possible images acquired using each microscope technique. It can be seen that almost all microscope techniques proved suitable to visualize individual bacteria that were clearly separated from each other or located towards the periphery of the tightly packed intracellular microcolonies. However, while many bacteria could be resolved laterally using the confocal (Fig. 1A), the images were sharper with clearly improved resolution and contrast using the iSIM (Fig. 1B) and Airyscan (Fig. 1C), and clearest of all when using the 3D-SIM microscope (Fig. 1D). Moreover, whereas individual bacteria could clearly be resolved laterally using both 405 and 488 dyes by 3D-SIM, with the shorter wavelength 405 excitable dye showing superior xy resolution as expected (Fig 1D), the images of Dylight 405-labelled bacteria were of significantly lower quality on the iSIM, confocal, and Airyscan (data not shown). We did not consider it productive to pursue this line of investigation further (e.g., to perform additional measurements) on these much poorer quality images, hence only the AF488 data is shown for the confocal, iSIM and Airyscan (Figs. 1A-C).

**Figure 1.**
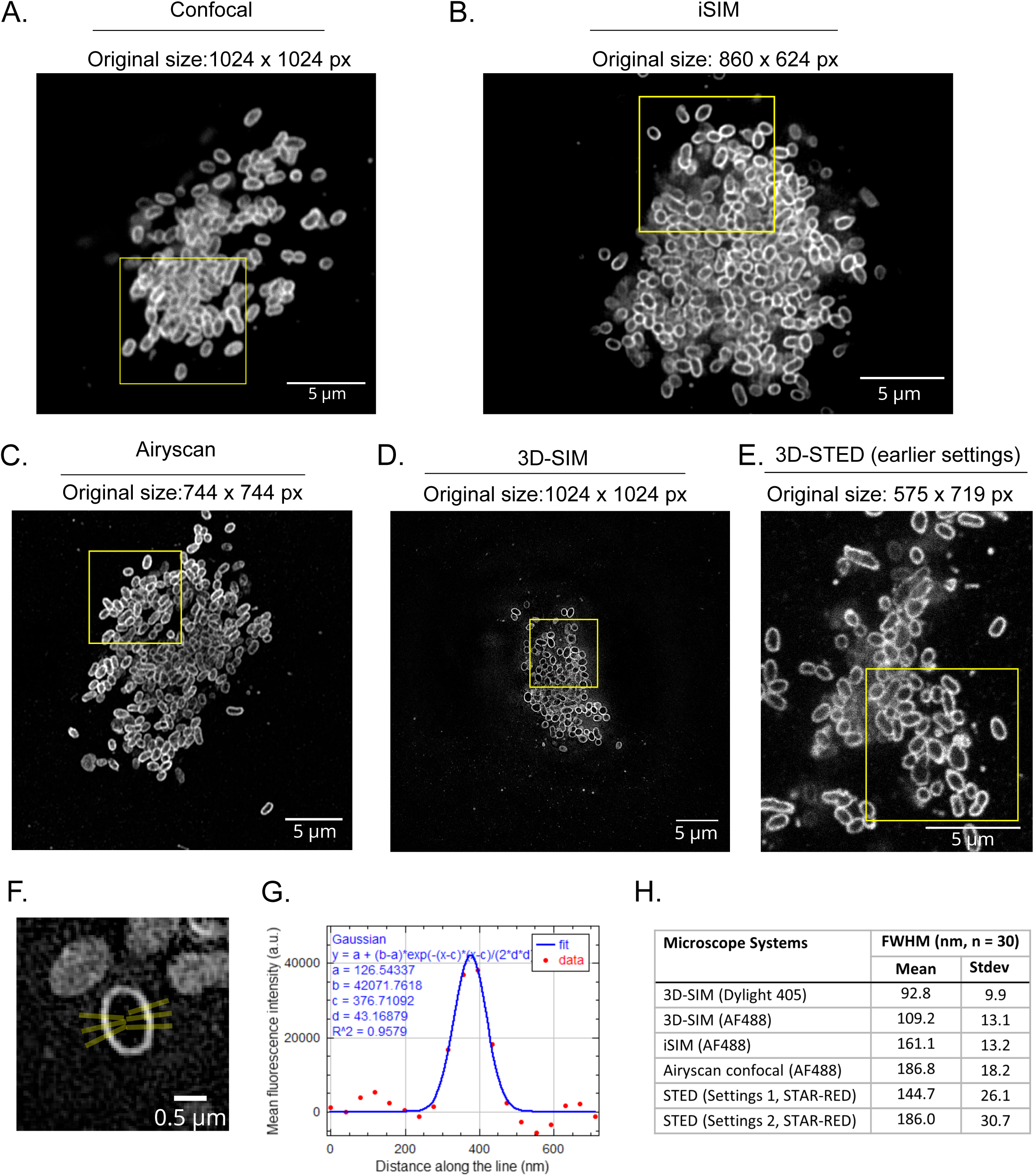
Comparison of microscopy techniques used here for imaging Ot bacteria. Lateral (XY) views of Ot bacteria aggregates inside HUVEC/HeLa cells stained with membrane marker ScaA using five different imaging techniques: (A) regular confocal, (B) iSIM, (C) Airyscan, (D) 3D-SIM and (E) 3D-STED. The yellow box in each image shows an 8x8 μm^2^ area and gives a perspective of imaging field size differences on different imaging platforms. (F) Cropped 3D-SIM (Dylight 405) image of a single Ot bacterium demonstrating how FWHM measurements were made. 6 line ROIs (3 on each side of minor axis, each 2 pixels wide) were drawn by the macro then adjusted to be as perpendicular to bacteria labelling as possible. (G) Example FWHM plot from one of line ROIs in Fig. 1F. In this plot, the red dots represent intensity measurements and the blue line is the fitting from built-in FIJI Gaussian fitter. Gaussian fitting formula, parameter values (a, b, c and d) and goodness of fit (R^2^) values are mentioned on the graph. (H) Mean FWHM derived from the different microscopy techniques: 3D-SIM (Dylight 405 and AF488), iSIM (AF488), Airyscan (AF488) and STED (both earlier and newer settings). For each technique the mean and standard deviation was calculated from 30 measurements.

For the highest quality image sets, we measured the full width half maximum (FWHM) distance as an indication of the relative resolution realised by each technique in practice (Fig 1H). Line ROIs (Fig 1F) were used to extract intensity profiles to which a Gaussian curve fitter was applied (Fig. 1G) to determine the FWHM. Further details are provided in the materials and methods section. ScaA is an integral outer membrane protein from the autotransporter protein family, and the anti-ScaA antibody interacts with an epitope on the extracellular domain of this protein^9^. Each labelled bacterium was encased by a single fluorescent line corresponding to surface exposed ScaA protein, thus the limiting factor on the measured thickness of this line was believed to be the microscope resolution, rather than the thickness of the biological structures themselves. FWHM analysis is often used as a parameter for comparing the resolution achieved by microscopes, though true “resolution” measurements are complex^35,47^. The maximum resolution achievable using a given microscope is often best illustrated by the separation of fluorescent markers spaced at known intervals on standardized samples, such as the Gattaquant nanorulers or Argolight slides^48,49^, however, this can still far exceed the practical resolution obtained on an actual biological sample. Here, the highest lateral resolution was obtained with the 3D-SIM system, combined with the shortest wavelength (405) dye, and the average FWHM for this system – 92.8 nm for 405 and 109.2 nm for 488 – was approximately consistent with the literature (110nm laterally for “green” emission^50^) which reports this technique to give double the resolution in each axis compared to conventional widefield fluorescence, leading to an approx. 8-fold volumetric improvement (reviewed by Schermelleh *et al.*^34^). The FWHM measured on the other super-resolution instruments was significantly greater, and did not match the predicted capabilities of these techniques. These have been quoted as ∼1.7x resolution increase in all axes on the Airyscan (we measured 186.8 nm laterally), whilst on the iSIM final measurements (after deconvolution) have been reported of around 145 nm lateral (we measured 161.1 nm) and 350 nm axial^17,19,34,51^. However, the resolutions achieved here by these methods did surpass that of conventional microscopy, typically quoted as 200-250 nm (lateral axis) and 500-700 nm (axial) for a high numerical aperture (NA) objective and green-emitting dyes. It should be noted that whilst every effort was made to perform FWHM analysis objectively, the higher-than-expected values could partially reflect the averaging of measurements made truly perpendicular to the surface with those where the measurement line glanced the surface and thus the fluorescent signal appeared thicker than the actual resolution limit.

For the standard confocal images, the significantly higher cytoplasmic background caused poor Gaussian fitting (R^2^ value below 0.9) or enlarged FWHM measurements (Supplementary Fig. 1). Whilst this could be improved with deconvolution or background subtraction, it’s likely that this would positively impact the resolution and present an unrealistically poor expectation for standard confocal images of this type of sample, so the data is omitted.

It was notable that one technique which proved unable to resolve bacteria at all, even in sparse groups, was 2D-STED microscopy, in contrast to 3D-STED, which was to prove the optimal choice for the questions asked here (see below). 2D-STED images appeared to show a diffuse label all over the bacteria, rather than a clear localization at the periphery of each of the bacteria (see, for example, left hand panel of Fig. 2A). We tested combining different proportions of 2D-STED and 3D-STED in the imaging parameters (effect on PSF demonstrated in Fig 2B), and found that as the percentage of 3D-STED increased, the labelling of the bacterial peripheries became much clearer (Fig 2A-from left to right) and the ability to accurately visualise them in the axial (z) axis also improved (Fig. 2C, representing the orthogonal view along the yellow lines in Fig 2A). We therefore restricted further analysis to samples imaged using 100% or 95% 3D-STED, both of which proved effective in resolving individual bacteria.

**Figure 2.**
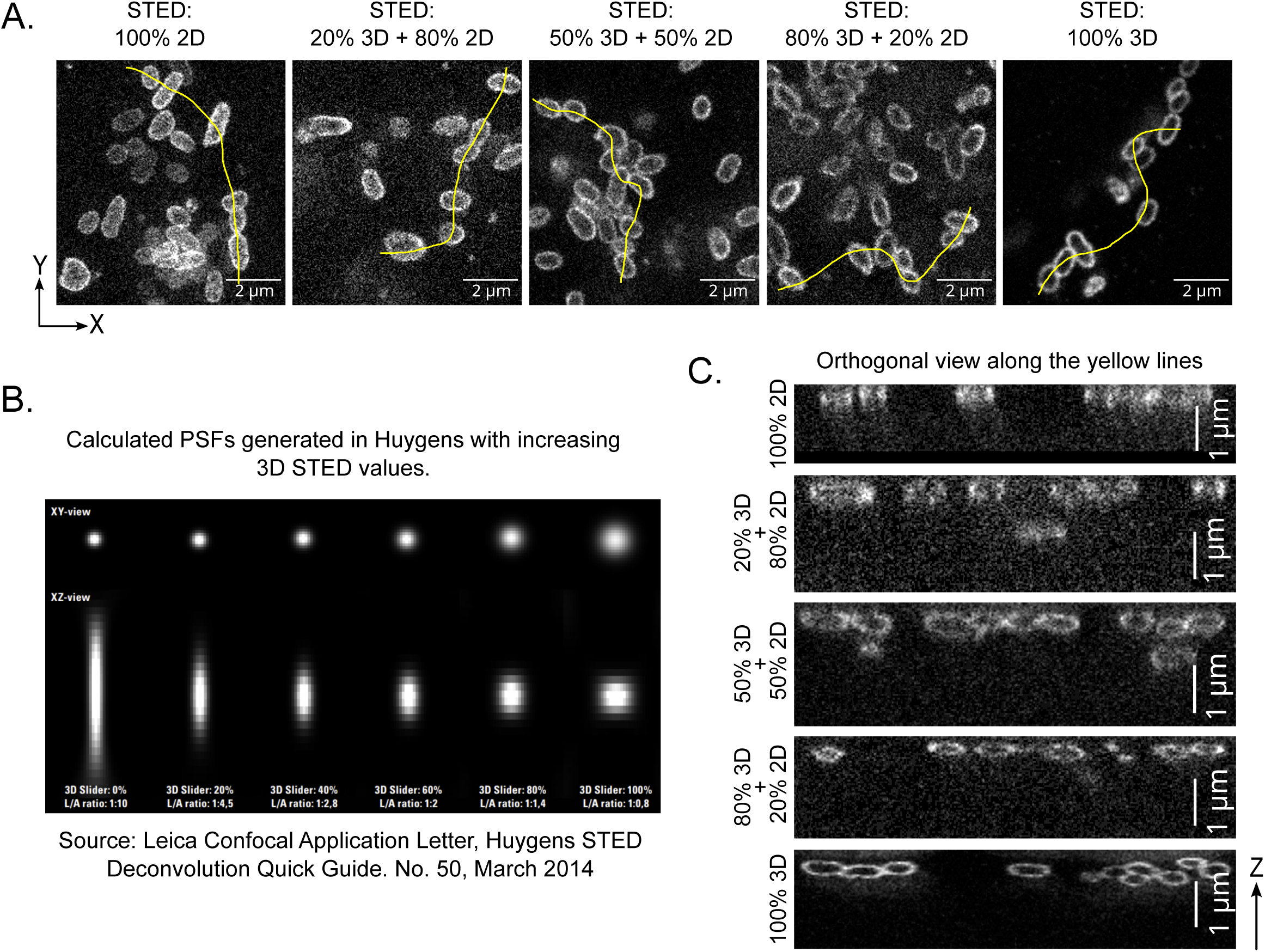
**Effect of combining different proportions of 2D-STED and 3D-STED PSF-shaping on the STED microscopy images.** (A) Lateral (XY) views of Ot bacteria images taken on the STED microscope using different percentages of 2D- and 3D-STED settings. (B) Illustration of the changes in lateral and axial PSFs (theoretical) as a function of different proportions of 2D- and 3D-STED, from 100% 2D on the left to 100% 3D on the right. (C) Orthogonal (XZ) views of Ot bacteria along the yellow line drawn in each panel of (A).

Ot has been shown to produce outer membrane vesicles that have been observed using transmission electron microscopy to be 50-150 nm in diameter^52^. To our knowledge these have not been reported using light microscopy. In our microscopy analysis we clearly observed ScaA-labelled ring-shaped vesicles budding off (Figs. 3Ai, Bi) and secreted close (Figs. 3Aii, Bii) to the surface of intracellular Ot bacteria when using 3D-SIM imaging at both 405 (Fig. 3A) and 488 nm (Fig. 3B), giving a diameter of ∼150-200 nm (Figs. 3Aiii, Biii), which is close to the reported vesicle sizes. Small spots, most likely corresponding to secreted vesicles, were also detectable in confocal, Airyscan and iSIM microscopy images (e.g. Fig 3Ci and ii), but they could not be resolved as true ring-shaped vesicles.

**Figure 3.**
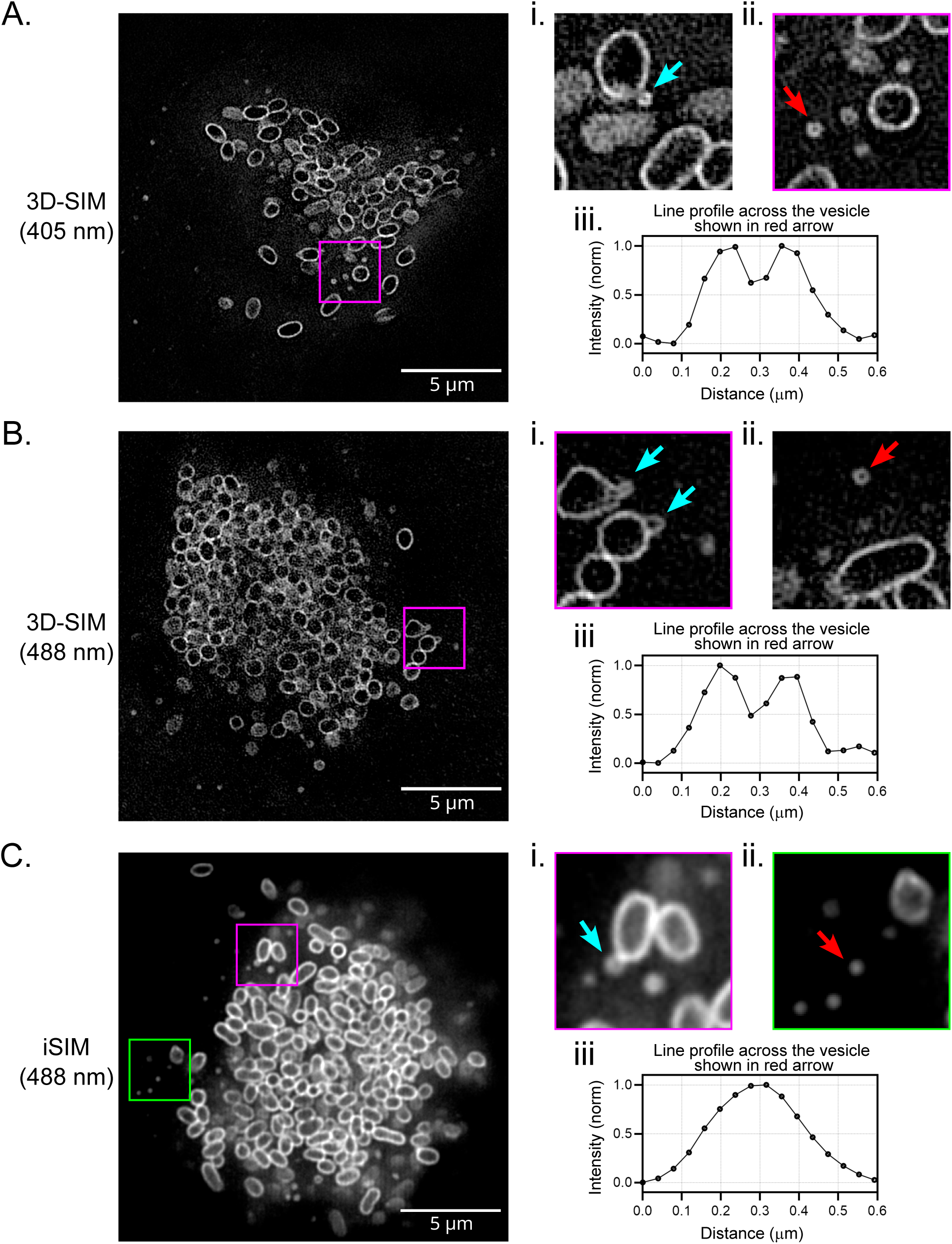
Characterization of Ot outer membrane vesicles. Lateral (XY) views of Ot bacteria images containing some examples of membrane vesicles using 3D-SIM imaging at wavelengths 405 nm (A) and 488 nm (B); and iSIM imaging at wavelength 488 nm (C). Panels in (i) and (ii) show zoomed in views of vesicles for the corresponding imaging technique. Panels in (i) show examples of vesicles budding off from the surface of Ot bacteria (cyan arrows). Panels in (ii) show examples of secreted vesicles close to Ot bacteria (red arrows). Graphs (iii) beneath the panels show line profiles across the vesicles identified by the red arrows.

### Only 3D-STED enabled clear delineation of individual bacterial cells throughout a tightly packed microcolony

One of the challenges associated with imaging intracellular Ot bacteria is that it is difficult to resolve individual bacteria within large three-dimensional tightly packed groups of cells. We therefore compared the ability of different microscopes to resolve individual bacterial cells within packed microcolonies. This required sufficient resolution in both the axial (z) and lateral (xy) axes, such that the limitations of certain super-resolution techniques became evident. Indeed, preliminary data with an external collaborator who specializes in the use of STORM microscopy had quickly revealed that this technique, based around a Total Reflection Interference Microscope (TIRF), proved completely unsuitable for resolving bacteria within intracellular clumps deep inside the cells (data not shown). We found that 3D-SIM, iSIM, confocal and Airyscan systems were unable to resolve all of the bacteria deep within clumps, though 3D-SIM could clearly resolve individual bacteria at the edges of infected cells or clumps (Fig 4). Only 3D-STED proved capable of resolving the outlines of all individual bacteria within the aggregates in the infected host cells (Fig. 4A-F).

**Figure 4.**
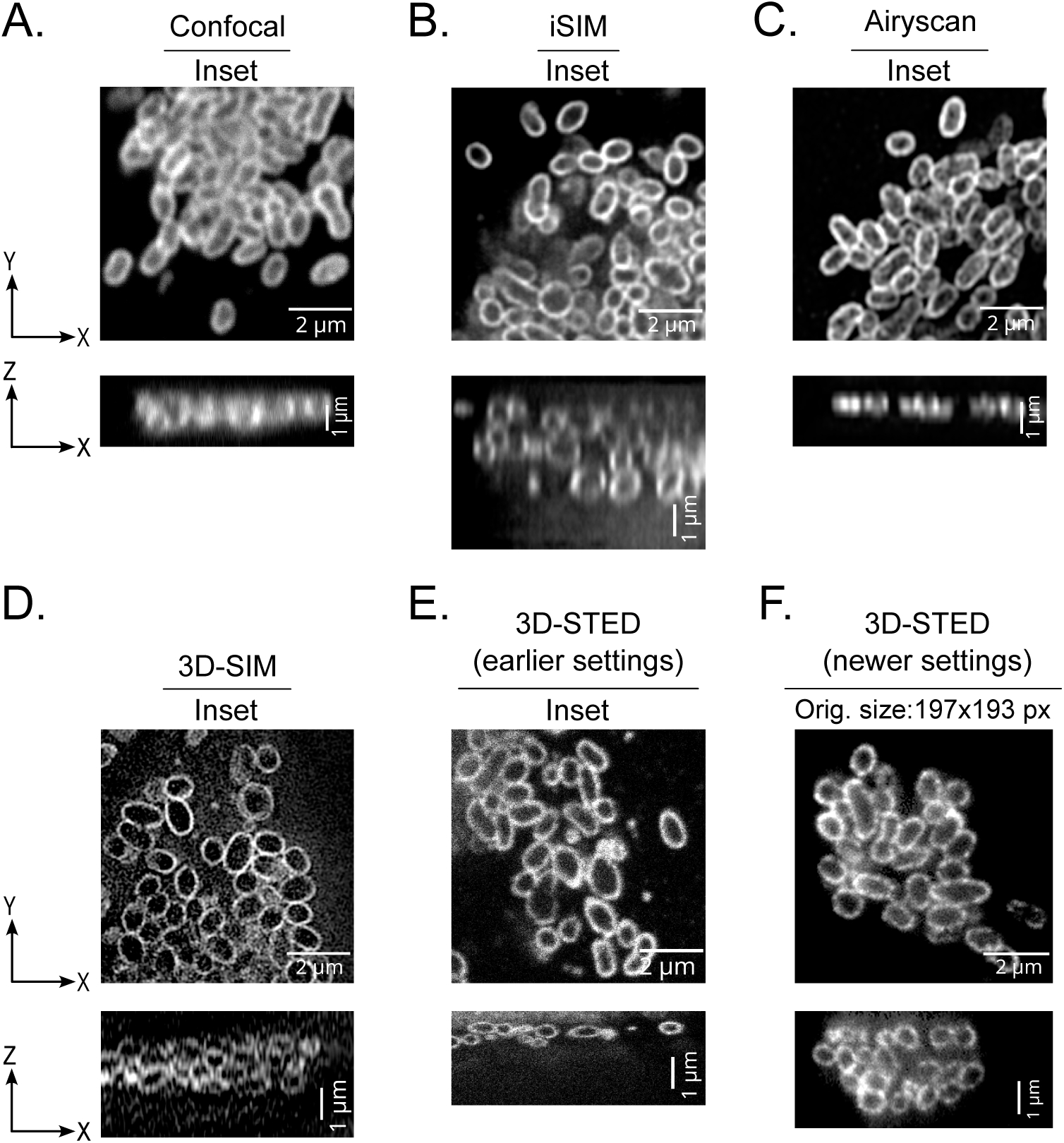
Comparison of super-resolution microscopy techniques for imaging Ot bacteria using lateral and axial views. Zoomed in lateral (XY) and axial (XZ) views of Ot bacteria aggregates inside cells using different imaging techniques and settings. The regions correspond to the yellow boxes in panels depicted in Figure 1A-E. Additionally, 3D-STED images with newer settings are also shown (F).

### Automated image segmentation of STED images reveals differences in bacterial morphology when grown in different cell types

Classical image segmentation techniques have been used for decades for the automated segmentation of cellular structures (whole cells, nuclei, membrane, subcellular organelles) in 2D images, but this process has always been onerous, if not impossible, for more challenging images of cells in a cluster and/or with low signal:noise such as the bacterial cells in the present work. Whilst capable bacterial segmentation pipelines, such as MicrobeTracker, allow individual cells to be automatically segmented and analysed^53^, their use is limited to analysing regularly shaped bacterial species arranged in two dimensions on a surface such as *E. coli* or *C. crescentus* grown on agar pads. Further, to analyse 3D datasets the complexity typically increases manyfold. To our knowledge, automated image segmentation and analysis have not previously been demonstrated for Ot. Here, based on the reproducibly high axial (z) resolution, as well as the clarity of imaging individual bacteria within large aggregates, 3D-STED images were used for developing a 3D segmentation and analysis pipeline.

There has been a recent surge in artificial intelligence-based methods, particularly deep learning methods for image segmentation^54^. Unlike classical methods, deep learning methods learn about the imperfections present in the images and after sufficient training often produce human-level segmentation accuracy for the structure of interest. For bacterial cells with membrane staining, a deep learning segmentation method called DeepBacs was published^55^, and it was demonstrated to work with 2D images. However, it is not clear if DeepBacs also works with 3D datasets of bacterial images. Another deep learning method for general cellular segmentation called Cellpose^56^, as applied here for 3D segmentation of Ot, is becoming popular amongst biological researchers and has been used for segmenting a wide variety of cell types and cellular structures^57^. We found that Cellpose was able to do accurate instance segmentation of bacterial cells in 3D-STED images, even for cells in aggregates (Fig 5A, Supplementary movie 1). We also tried Cellpose on iSIM (Fig. 5B) and (Fig. 5C) 3D-SIM images, but the segmentation results were sub-optimal, particularly in areas where cells were clustered together. Cellpose generates 3D cell masks by combining the 2D segmentation masks from images in all three orthogonal directions - XY, YZ and XZ. Even though all three imaging techniques (3D-STED, iSIM and 3D-SIM) produced good XY resolution images in thin layers or at the periphery of clumps, the resolution in orthogonal directions (YZ and XZ) was significantly worse for iSIM and 3D-SIM than when using 3D-STED (Figs. 5A-C), leading to worse 3D cell segmentation in iSIM and 3D-SIM images, but accurate 3D cell segmentation in 3D-STED images. Next, we imported Cellpose-generated segmentation masks for 3D-STED images into Imaris software and quantified 3D cellular parameters such as volume, sphericity and ellipticity of Ot cells. We tested the ability of this pipeline to resolve biologically meaningful differences in bacterial morphology by comparing bacterial dimensions when grown in two different cell lines. We found that the volume of Ot bacteria does not significantly change whether they are inside HeLa cells or in HUVEC cells (Fig 6A), but their shape is significantly different in these two cell lines. The Ot bacteria were found to be more elongated in HUVEC cells than in HeLa cells, as shown by both sphericity and ellipticity measurements (Figs 6B, C).

**Figure 5.**
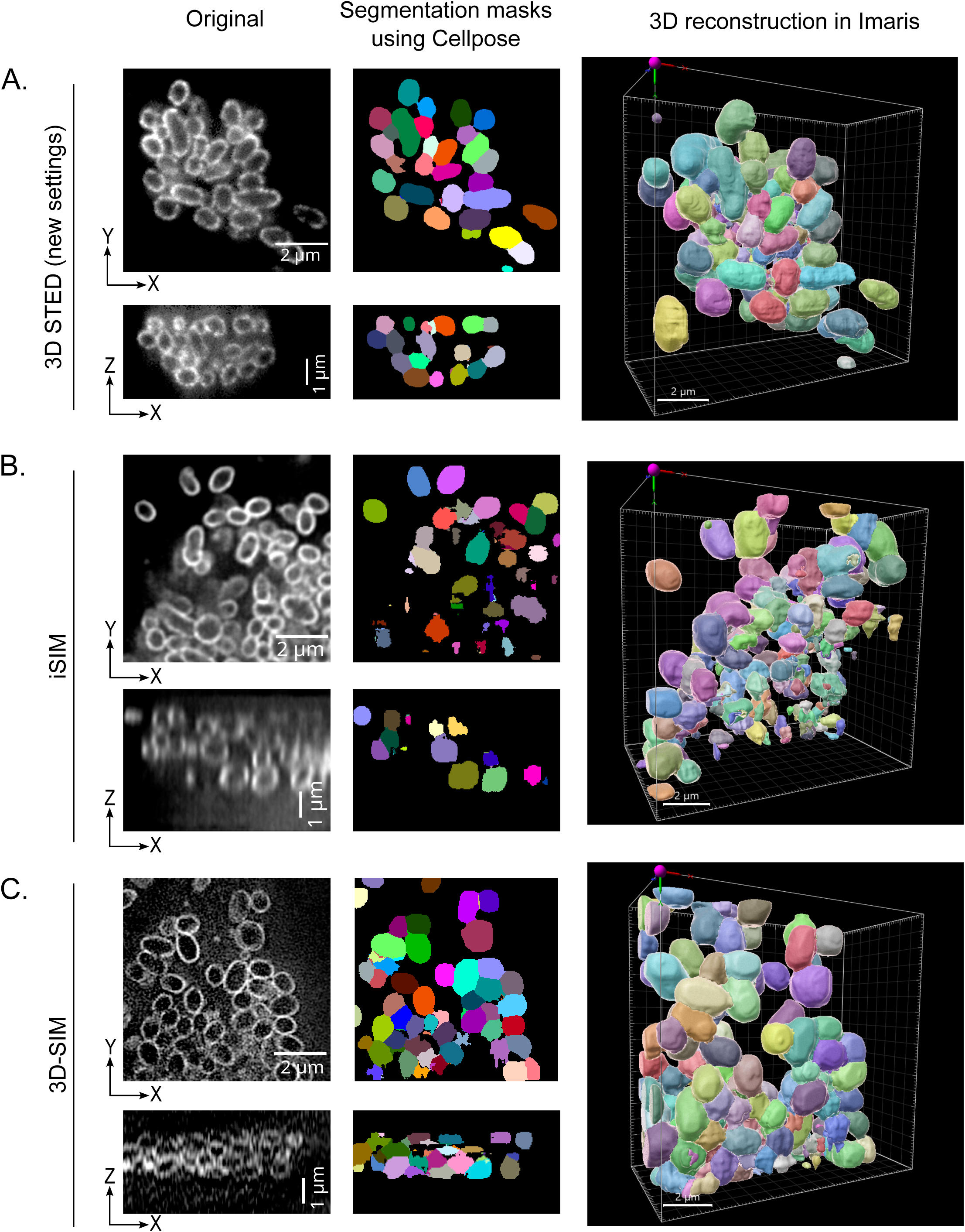
The deep learning-based 3D segmentation and reconstruction analysis comparing three different imaging techniques. Left most images show lateral and axial views of 3D-STED (A), iSIM (B) and 3D-SIM (C). Middle panels show the corresponding segmentation masks generated with Cellpose. Rightmost panels show 3D reconstruction of individual bacteria using segmentation masks in Imaris.

**Figure 6.**
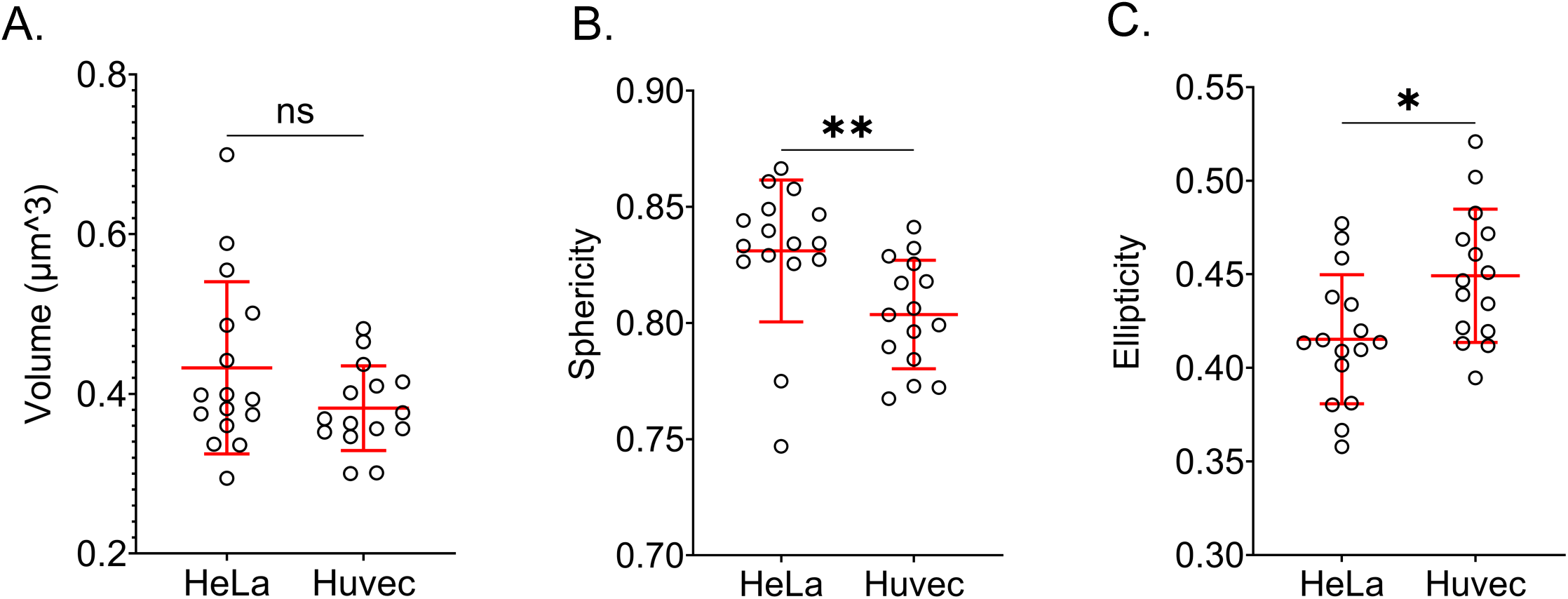
Comparison of Ot bacteria size and shape parameters when grown in HeLa vs HUVEC cells. Quantification of Ot bacterium volume (A), sphericity (B) and ellipticity (C) using best overall (XYZ) resolution 3D-STED images. Data shows elongated Ot bacterium morphology in HUVEC cells compared to in HeLa cells. Each point represents an average measurement value for one image containing ∼70 bacteria. In total, n=1119 (HeLa) and 1024 (HUVEC) Ot bacteria were analyzed; unpaired two-tailed Mann-Whitney test, ns = 0.28 (A), **p = 0.0015 (B) and *p = 0.015 (C).

## DISCUSSION

### The most suitable microscope technique to be used depends on the nature of the question being asked and the size and structural organization of the biological specimen

The microscopy technique selected for any given research project will need to be determined by the exact biological question. If all that were asked here was whether the bacteria were growing successfully in these cell lines, then a simple widefield microscopy approach would suffice, but for imaging and characterising individual cells within the tightly packed intracellular microcolonies of Ot, confocal microscopy had already proven inadequate^9,58,59^. Many reviews have already addressed the advantages and disadvantages of different super-resolution modalities in general and for answering specific questions: factors to be taken into consideration include the exact resolution required, the number of different labels that must be used together, the thickness of the biological specimen and the speed of imaging required, which can be crucial if the cells are live^17,33–35^. Some practical comparisons of different modalities have been reported^50^. However, few core facilities, let alone individual laboratories, are fortunate to have a comprehensive range of super-resolution techniques available to compare on equivalent samples prepared by the same researchers and imaged by one team at the same time and location, thereby minimising reproducibility issues. We initially expected this to be a fairly straightforward biological specimen to image by any of the super-resolution approaches featured here, given the requirement to image only at the single bacterial cell level, rather than subcellularly, the need to use only one colour and because we were working with fixed cells. To our surprise, however, only a single modality – 3D-STED - proved capable of addressing all questions asked in this project. The reason for this lies in the small dimensions and tight packing of these bacteria in their intracellular colonies, and the consequent need for high axial resolution across several micrometers of depth, in addition to sufficient lateral resolution. This was clearly illustrated by the inability of 2D-STED – the mode which would offer the highest lateral resolution on this instrument – to cleanly resolve the bacterial peripheries. We surmise that for 2D-STED, where the axial dimension of the PSF is not improved over widefield mode (i.e. is > 500 nm), every voxel included signal from the surface of the bacteria either above or below the plane of focus, resulting in an apparently diffuse fluorescence over the whole of each bacterium. As we increased the proportion of 3D-shaping of the PSF, which is accomplished easily on the Abberior Facility Line instrument by way of a simple slider, its axial dimension was reduced to below the thickness of the bacterial cells. Thus with 95% or 100% 3D-STED, where an isotropic resolution of around 80-90 nm is attainable on commercial systems^19^, we could image just the medial section of each bacterium, thus capturing signal only from the cell surfaces at the periphery.

3D-SIM, using a short wavelength (blue-emitting) dye, gave excellent resolution in the lateral (xy) axis, and enabled all bacteria but those in the centre of the clumps to be readily resolved and counted. For thin layers of bacteria, rather than large intracellular clumps, we would thus recommend this technique as the best modality, being quick to use and, in our hands, giving highly reproducible super-resolution results. 3D-SIM, however, relies on the propagation of the patterned illumination through the entire thickness of the sample. Spherical aberrations, a particular problem with thicker samples with varying refractive indices, reduce the amplitude and transfer of high spatial frequency information, leading to poor image reconstructions with significant aberrations^37^. Moreover, the axial (x) resolution of this system is at best >250 nm, even with blue-emitting dyes, hence the default optical section spacing is set by the system to 125 nm (to satisfy Nyquist sampling). Together, these factors prevented the successful resolution of individual bacteria at the interior of the aggregates and 3D-SIM thus proved insufficient for successful image segmentation and 3D measurements.

The iSIM and Airyscan methods also proved superior to standard confocal microscopy in resolving thin layers of bacteria in two dimensions. These techniques are typically better suited than 3D-SIM towards thicker samples, due to the rejection of background fluorescence by pinhole optics prior to image formation^17^, but their inability to match the axial resolution of the 3D-STED still limited their usefulness for these specific samples.

### Commercial super-resolution instruments require advanced understanding and systematic testing to ensure optimal performance

Over the past decade, the number and range of commercially available super-resolution instruments has increased enormously^17,33^. The production of some have even been discontinued recently, including, to our great disappointment, the OMX 3D-SIM instrument used in these studies. Increasingly, these instruments are marketed as “easy-to-use” for the average biological researcher, and it is common for the companies to claim a maximum achievable resolution for their instrument. But is this truly achievable in practice, by biologists and on real biological samples? The work presented here was undertaken in a well-staffed core facility with significant experience running super-resolution microscopy experiments on a variety of samples. All the instruments used were installed at least 2 years prior to this study, some considerably more. Yet, many lessons arose from this work, and the results were often surprising, hence there is great value in sharing the outcomes in order to reduce the ramp-up time for others starting a similar study.

Firstly, it is unclear to us why only the OMX system, using an Evident (Olympus) objective, proved compatible with 405-labelled samples. Presumably this reflects a problem in the hardware specifications in each case for the confocal, Airyscan and iSIM systems, whether it was suboptimal transmission and/or reflection of the shorter wavelengths by the objective lens or other optics internal to the system, or a problem of system alignment in this channel. Whatever the cause, this could be another reason why few publications appear to take advantage of the shorter wavelength, higher resolution blue-emitting dyes.

Secondly, it is very difficult to acquire consistent and reproducible data over a significant time period, because the instruments themselves, as well as their operation, are constantly evolving. In the case of the Abberior STED in particular, the Facility Line system is still relatively early in its development, and has been evolving rapidly since it was installed in the Rockefeller University’s Bio-Imaging Resource Center (RU BIRC). During that time we have switched to a new principal objective lens (100x/1.45 XApo objective, instead of 100x/1.40 SApo objective), a Deformable Mirror for Adaptive Optics has been installed, and there have been many updates to the software. Moreover, the service on this system switched from largely remote (during the COVID pandemic) to mainly in-person visits, which can only have been beneficial for the alignment of this complex system. And perhaps the most influential factor of all, the staff operating this system gained enormous experience in how best to tune and optimise the acquisition settings. This is the main reason for such different settings between the first set of STED experiments, generating the images used for the FWHM measurements (Figs. 1 and 2, and Fig. 4E), and the latter experiments, investigating the shape of the Ot bacteria when grown in different human cell lines (see Fig. 4F). For the latter study, for example, the depletion laser power was set much lower, while the pixel size was increased from 25 to 40 nm (in x,y and z) (both of which would reduce photobleaching but also, potentially, resolution), partly because it had now been established that this was still sufficient to resolve the bacteria in 3D, and partly because the imaging was optimized by the implementation of Adaptive Optics and Adaptive Illumination modalities^60^ (which reduce bleaching and thus increase the potential for improving resolution). Operator experience in choosing the most appropriate acquisition settings and in learning how to check the system for optimal alignment, and, importantly, in working alongside the company to attain it, only comes from hours/days/months of work on a given system to become truly familiar with its intricacies. This is why such instruments are generally best placed in a core facility setting where the considerable experience of the staff is available to each user, and we urge researchers to consult their core facility experts, including both microscopy and analysis experts, as early as possible in experimental conceptualisation and design to save a lot of potential wasted time, effort and reagents.

It is also critical to have an accurate and reproducible method of measuring the true resolution of each instrument on a consistent basis. Subresolution fluorescent microspheres (beads) are typically used for such measurements, but most preparations of these are 2-dimensional and thus do not allow measurements of axial (z) resolution. For this, more recent commercial standard slides such as the Argolight slide^49^ provide new opportunities but are not applicable to systems such as STED and STORM where the resolution achieved depends on the use of specific dyes^47^.

Furthermore, the resolution deeper into a biological sample is significantly deteriorated by factors such as refractive index mismatch between the immersion medium and the specimen. The work presented here illustrates the critical nature of international community endeavours, such as those of QUAREP-LiMi^61^, to establish common standards for measuring and reporting microscope performance across time and on different samples. An understanding of the common technical problems such as suboptimal objectives or system misalignment is especially important for the more “workhorse” microscopes such as the confocal system used here, which is used in hundreds or thousands of laboratories worldwide by less experienced operators who may be entirely unaware of the potential for performance drifts over time.

### Could we have achieved even higher resolutions using deconvolution or recent methods combining deep learning with super-resolution during acquisition?

It is important to note that we are not using the FWHM measurements reported in this study to claim that this is the maximum resolution achievable on each system. First, we could have improved the resolution by optimizing the sample labelling further, for instance by using directly labelled antibodies or nanobodies^33^. Second, we could probably have improved the STED resolution even further, by more stringent optimization of the depletion laser power and by applying the adaptive illumination methods, ResCUE and DyMin^60^, used for the later bacterial segmentation pipeline (Fig. 4F) that allow you to minimize photobleaching and thus improve signal-to-noise by only employing the depletion laser where signal is demonstrated to be present. We tested out all of these methods, but for the final experiments we opted for the least stringent acquisition settings that were sufficient for analysing bacterial shape.

Third, for this study we chose to use the image processing steps that were recommended by the companies for each commercial system tested. In the case of the iSIM, the use of deconvolution software was recommended as the second step to achieve the predicted final resolution for this system, and therefore it was applied here. We regularly use other deconvolution software packages, such as SVI Huygens, to improve both resolution and, more significantly, contrast in images. Still, the debate over whether deconvolution methods should be applied to all images remains contentious and we did not want to detract from the takeaway message of this story. Since the raw 3D-STED data provided sufficient resolution to segment the bacteria within aggregates inside cells, there was no requirement to apply an additional step of deconvolution here. Nevertheless, for studies where the raw data is unsuitable or inadequate to perform the required measurements, we recommend applying quantitative and well-documented deconvolution methods with a clear methodological declaration.

A recent publication demonstrated improved axial resolution by combining deep learning approaches with 3D-SIM to achieve isotropic ∼120 nm resolution on labelled bacteria, enabling the visualization of components inside bacteria^62^. This approach provides the advantage of being applicable to other microscopy methods, perhaps even confocal microscopy, which would greatly expand the availability of imaging equipment suitable for microbiologists working at the intracellular level. However, the development of such multi-step deep learning pipelines is time-consuming, and the success of this study also arose from the extreme care and attention to detail that the authors paid to refractive index matching and sample preparation. Moreover, researchers applying Artificial Intelligence (AI) methods to enhance the resolution of their images need to be aware of the potential for introducing artifacts^35^. For studies requiring extended timelapse imaging of living cells, this approach could well prove superior to the photobleaching-prone 3D-STED approach used here, yet the method we used here was much quicker and simpler and proved sufficient to answer our biological question.

### Is Cellpose the best deep learning method for 3D segmentation of Ot bacteria?

In this study, we tried several popular deep learning methods for 3D segmentation of bacterial cell, such as StarDist^63^, PlantSeg^64^, DeepBacs^55^ and Cellpose^56^. We found that Cellpose worked the best for Ot bacteria images acquired by 3D-STED microscopy. Cellpose did not perform well on Ot bacteria images acquired on iSIM or 3D-SIM microscopes, where the Z-resolution was poor compared to what we could obtain by 3D-STED. One caveat with these results is that Cellpose could potentially be used to train new deep learning models for iSIM and 3D-SIM images, which might lead to segmentation accuracy comparable to what is currently obtained using 3D STED. Similarly, any of the other deep learning methods tested (StarDist, PlantSeg, DeepBacs) could be trained on our Ot bacteria images to possibly reach the segmentation accuracy of Cellpose. We chose not to train a new deep learning model for this study, as this is a very resource-intensive process and might not be feasible for many biological researchers. Here, our goal was to evaluate the robustness of already trained deep learning methods for 3D bacterial segmentation and we demonstrated that Cellpose can be successfully used for 3D segmentation of Ot bacteria in images acquired on a near isotropic resolution microscope such as 3D-STED.

In summary, we demonstrate that there are distinct advantages and disadvantages of using various super-resolution imaging platforms to image intracellular Ot and reinforce the view that the appropriate microscope should be selected based upon the biological sample and the specific question being addressed. However, two new analyses arising from our study stand out for their potential impact on the obligate intracellular bacteria field. First, the ability to resolve and quantify bacterial morphology using STED will enable studies into Ot cell growth and division and how this is affected by antibiotics, the bacterial stage of development, and interactions with host cell pathways. Second, the ability to directly image outer membrane vesicles of Ot will support studies into the synthesis and function of these poorly understood cellular structures. Together these imaging platforms have the potential to yield novel insights into the biology of this important but poorly studied human pathogen, as well as other organisms presenting similar imaging challenges.

## MATERIALS AND METHODS

### Bacteria strain, cell line, and culture conditions

The clinical isolate *O. tsutsugamushi* strain UT76 was used throughout the experiments^65,66^. Bacteria were grown in either human cervical epithelial HeLa cells (ATCC CCL-2) or primary Human Umbilical Vein Endothelial Cells (HUVEC) (ATCC PCS-100-010). HeLa cells were cultured in DMEM (41965-039, Gibco) supplemented with 10% FBS (16140-071, Gibco). HUVEC cells were cultured in human large vessel endothelial cell basal medium (M200500, Gibco) supplemented with 10% LVES (A1460801, Gibco). All cells were maintained at 37°C and 5% CO_2_ atmosphere. For bacterial propagation, bacteria were grown in mouse fibroblast L929 cell (ATCC CCL-1), as described previously^67^.

### Sample preparation for microscopy

Coverslips (High precision No 1.5H, Paul Marienfeld GmbH) were pre-sterilized by incubating with pure ethanol for 15 min followed by exposure to UV light for 30 min. Sterile coverslips were then pre-coated with 5ug/ml fibronectin in sterile PBS (pH 7.4) for 30 min. HeLa cells (2 × 10^4^) or HUVEC cells (4 × 10^4^) were seeded onto pre-coated coverslips and incubated overnight. Cells were infected with frozen stock UT76 (∼MOI 1:100) for 4 days before fixation with 1% formaldehyde in PBS for 15 min at room temperature. Cells were permeabilized by incubation in absolute ethanol for 1 h on ice followed by 0.5% Triton-X100 in PBS for 30 min on ice. Bacteria were labelled with an anti-ScaA antibody (AP40436, custom made by abclonal)^9^ at a dilution of 1:200 in PBS buffer containing 1mg/ml BSA for 1 h at 37°C. Cells were washed twice with PBS-BSA and were then incubated with secondary antibodies (dilution 1:1000) appropriate for each microscope at for 30 min at 37°C in the dark together with Hoechst (Molecular Probes, Hoechst 33342, trihydrochloride, trihydrate; dilution 1:1000 in PBS) to label host cell nuclei. Secondary antibodies used were as follows: goat anti-rabbit STAR-RED (Abberior GmbH 90725CW-5); goat anti-rabbit AlexaFluor 488 (Thermo Scientific A11008); goat anti-rabbit AlexaFluor 594 (Thermo Scientific A11012); goat anti-rabbit DyLight 405 (Thermofisher 35551). Coverslips were washed with PBS 3 times, mounted on glass slides using uncured Prolong Diamond Antifade Mountant (P36961, Invitrogen/ThermoFisher), and sealed around the edges with quick-drying nail polish.

### Microscopy 3D-SIM

3D-SIM super-resolution microscopy data were acquired using a DeltaVision OMX V4/Blaze 3D-SIM super-resolution microscope (GE Healthcare/Cytiva/Leica Microsystems). This OMX system was fitted with a 100x/1.40 NA UPLSAPO oil objective (Olympus/Evident), three Evolve electron-multiplying charge-coupled device (EMCCD) cameras (Photometrics/Teledyne) that were used in EM gain mode set at 170 electrons/count; 405nm or 488 nm laser lines for excitation, a Sedat Quad dichroic and 436/31 or 528/48 emission filters, respectively. Optical sections were acquired at 125-nm intervals. Immersion oil refractive index (R.I.) was selected to optimize for each channel and the ambient temperature inside the cabinet. Structured illumination data sets were reconstructed using softWoRx 7.2.2 software (GE Healthcare/Cytiva/Leica), employing optical transfer functions (OTFs) generated from empirically measured point spread functions acquired from 100 nm yellow-green FluoSpheres (505/515, F8803 Invitrogen/Thermofisher) or 170 nm blue FluoSpheres (360/440) (P7220 Invitrogen/ThermoFisher). Since single colour labelling was used for each slide, the OTFs were acquired using oil whose R.I. was optimized for each channel, according to procedures described by Demmerle et al.^37^. Structured illumination data sets were reconstructed using channel-specific k0 values and a Wiener filter of 0.001. The effective pixel size of the reconstructed 3D-SIM images is 40 nm in xy and 125 nm in z.

### iSIM

Images were acquired on a VisiTech International instantSIM (VT-iSIM) system, mounted on a Leica DMi8 stand fitted with a 100x/1.40 NA HC PLAPO CS2 immersion objective lens, and controlled using VisiView 4.5.0.13 software Visitron Systems GmbH). DyLight 405 labelled bacteria were illuminated using a 405 nm laser line, a Quad zt405-488-561-640rpc polychroic (Chroma) and 450/50 Emission filter (Chroma). Alexa 488 labeled bacteria were imaged using a 488 nm laser line, a Quad zt405-488-561-640rpc polychroic (Chroma) and 525/50 Emission filter (Chroma). A tube lens of 2.0x magnification was used for the 488-labelled samples to better match the 6.5 micron pixel size of the Hamamatsu ORCA Fusion Gen-III camera), resulting in an effective pixel size of 32.5 nm on the image. This extra 2.0x magnification could not be used for the 405 labelled samples, as the emitted light intensity was already too low with a 1.0x tube lens to acquire high quality images. Microvolution deconvolution software version 2020.4.1.0, installed on the system and employed via the VisiView interface as an integral step of image acquisition, was used to increase the resolution of the final images.

### Confocal

Confocal images were acquired on a Zeiss LSM880 on an AxioObserver stand fitted with a 63x/1.40 PlanApochromat Oil DIC M27 lens. The Main Beam Splitter was (MBS) 488/561/633, the Ex wavelength was 488 nm and the emission filter set to 505-565 nm. The software used for the acquisition was Zen Black 2.3. The sample was scanned unidirectionally with 2x averaging and a pixel dwell time of 0.78μs. The signal was detected with a GaAsP detector with the Gain set at 600 and the detection wavelength range set at 500-565nm. The images were acquired as a standard 1024x1024 size image which, depending on the zoom used, resulted in a variable pixel size in xy, and in some of the images it met Nyquist sampling, and in others resulted in oversampling. All images were acquired so that the z-step of the z-stack met Nyquist sampling.

### Airyscan confocal

Images were acquired on a Zeiss LSM880 with an AxioObserver stand fitted with an Airyscan1 and a 63x/1.40 PlanApochromat Oil DIC M27 lens. The Main Beam Splitter was (MBS) 488/561/633 and the excitation laser used was 488 nm. The sample was scanned using Zen Black 2.3 in Airyscan SR mode, unidirectionally, and without averaging. The signal was acquired on the Airyscan1 detector with the gain set at 750 and the emission filter was 495-550 nm. The images were acquired using the optimal settings for sampling as was determined by the acquisition software in order to satisfy Nyquist sampling in both xy and z. The resulting Airyscan images were processed with the Wiener filter set at auto strength.

### STED

STED images were acquired on an Abberior Instruments Facility Line system fitted with a 775 nm depletion laser and four APDs, mounted on an Olympus IX83 motorized stand fitted with a ZDC2 hardware autofocus system and controlled using LightBox software v. 16.3.14287-w2129. The first set of images were acquired using a 100x/1.40 UPLSAPO100XO (Olympus/Evident) oil immersion objective. The microscope system was adapted and improved over time, with the addition of a 100x/1.45 NA UPLXAPO100X (Olympus/Evident) oil immersion objective, and a deformable mirror-based adaptive optics system (Abberior Instruments). In addition, the operators of the system gained experience and a deeper understanding of the optimal parameters for 3D-STED imaging as the research progressed. Thus, both microscope configuration and acquisition settings were different for the images acquired later for cell shape analysis. For both data sets, STAR-RED-labelled samples were excited using a pulsed 640 nm laser line and the emission range was set to 650-755 nm, and AlexaFluor 594-labelled samples were excited using a pulsed 561 nm laser line and the emission range of 588-698 nm. In both cases the STED setting was optimized at 95% 3D-STED; the pinhole was set to 0.72 Airy Units; time gating and adaptive illumination were engaged and optimized (ResCue and DyMIN); and the pixel size set to be the same for all 3 axes (X,Y,Z). In the latter images for cell shape analysis (Fig. 4F), the Adaptive Optics setting was also engaged.

### Full Width Half Maximum (FWHM) measurement for relative resolution performance comparison

We compared the relative lateral (xy) resolution performance of each system using images of bacterial cells in intracellular clumps, by taking FWHM measurements of labelled ScaA using two macros written in the open-source image processing software, ImageJ/Fiji^68^. The first macro created 3 line Regions of Interest (ROIs; 4 pixels apart) on each side of the bacterial cell. Manual adjustments were made to ensure the line ROIs were as perpendicular to cell wall as possible (Fig. 1F). The second macro was used to loop through each line ROI and from each extract an intensity profile (2 pixels wide), then run the ImageJ built-in Gaussian curve fitter (Fig. 1G) to determine sigma (σ), the standard deviation of the Gaussian function, and to calculate the Gaussian FWHM using equation, Gaussian 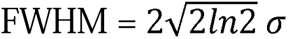.

To ensure an unbiased robust approach, we limited ourselves to use only unprocessed images of bacteria located within 2 microns of the coverslip surface. As above, FWHM measurements were taken at 6 locations for each bacterium (3 line ROIs on both minor axis sides of bacterium; Fig. 1F). Fig. 1G shows one of the intensity profiles (red dots) with Gaussian fitting (blue line) from a line ROI in Fig. 1F. For each microscope and labelling technique, 30 FWHM measurements were taken from 5 bacteria exhibiting good separation from neighbouring bacteria. Then the mean FWHM and standard deviation were calculated (Fig. 1H).

### Pipeline for the Ot cell segmentation, counting and cell shape analysis

HeLa or HUVEC cells infected with the ScaA-labelled Ot bacteria were imaged on different microscopes. 3D-STED images were acquired isotropically (voxel size: 40nm x 40nm x 40nm), whereas images from other techniques were acquired using Z-step sizes recommended by the acquisition software to ensure Nyquist sampling. For 3D segmentation, non-isotropically imaged volumes were converted to isotropic volume using BioVoxxel Toolbox^69^ in Fiji/ImageJ 2.1.4.0/1.54i. The 3D Gaussian blur with radius=2 was applied to reduce the noise. Cells were segmented in Cellpose 2.0.4 using the *cyto2* model with a diameter of 40 pixels. The bacterial cell shape analysis was performed in Imaris 9.9.0. All bacteria labels touching the image boundary or with very small volume (< 0.15 μm^3^) were filtered out. Bacterial volume, sphericity and ellipticity (prolate) were calculated. The sphericity is defined as the ratio of the surface area of a sphere with the same volume as the given cell to the surface area of the cell 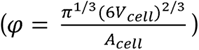. For ellipticity calculations, cells were modeled as American football shape (prolate ellipticity defined as 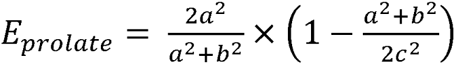, where a, b and c are the lengths of the three semi-axes of the ellipsoid). Data were exported from Imaris as CSV files, aggregated in Excel and plotted using GraphPad Prism 9.4.1.

### Generation of orthogonal views

The *Orthogonal Views* command in Fiji/ImageJ 2.1.4.0/1.54i was used to generate the orthogonal XZ view of the 3D Z-stack. The *Dynamic Reslice* command was used to generate the orthogonal view along manually drawn freehand line ROIs.

## ACKNOWLEDGEMENTS

AN, VS and CP are indebted to the Rockefeller University’s support of the Frits and Rita Markus Bio-Imaging Resource Center (RRID:SCR_017791). Ved Sharma is further supported by the Arnold and Mabel Beckman Foundation(dx.doi.org/10.13039/100000997) award ID: AWD00000407 for Light-Sheet microscopy and Data Science. The OMX 3D-SIM microscope in the BIRC was funded by Award Number S10RR031855 from the National Center for Research Resources. JLSY, GDW and the A*STAR Microscopy Platform (AMP) are funded by National Research Foundation (NRF) Shared Infrastructure Support grant for SingaScope (NRF2017-SISFP10) and by the Agency for Science, Technology and Research (A*STAR). JS was supported by a Wellcome Trust Senior Research Fellowship (224277/Z/21/Z) and an NIH R56 award (R56AI148645).

---

**Supplementary Figure 1.**
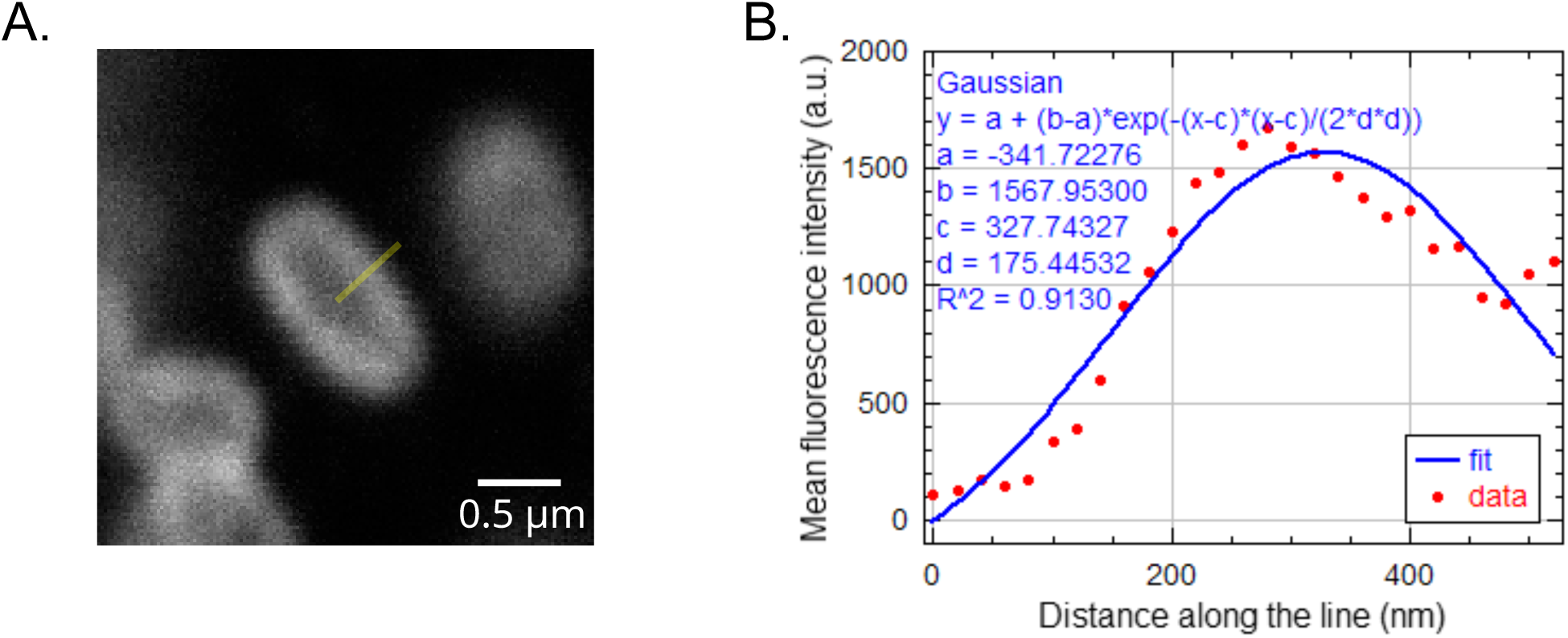
High cytoplasmic background in standard confocal image resulted in poor curve fitting, resulting in an enlarged and inaccurate FWHM measurement. (A) Cropped confocal image of Ot bacteria (AF488 labelling). Confocal images exhibited high cytoplasmic background inside the bacteria, much higher than the background outside the bacteria. (B) FWHM plot of line ROI. Gaussian fitting formula, parameter values (a, b, c and d) and goodness of fit (R^2^) values are mentioned on the graph. Due to the difference in background between inside and outside of the bacterium, the built-in FIJI Gaussian fitter exaggerated the fitting and this resulted in an inaccurate measurement of resolution.

## REFERENCES

1 den Blaauwen, T., Hamoen, L. W. & Levin, P. A. The divisome at 25: the road ahead.Curr Opin Microbiol 36, 85–94, doi:10.1016/j.mib.2017.01.007 (2017).

2 Rowlett, V. W. & Margolin, W. The bacterial divisome: ready for its close-up. Philos Trans R Soc Lond B Biol Sci 370, doi:10.1098/rstb.2015.0028 (2015).

3 Jones, L. J., Carballido-Lopez, R. & Errington, J. Control of cell shape in bacteria: helical, actin-like filaments in Bacillus subtilis. Cell 104, 913–922, doi:10.1016/s0092-8674(01)00287-2 (2001).

4 Ma, X., Ehrhardt, D. W. & Margolin, W. Colocalization of cell division proteins FtsZ and FtsA to cytoskeletal structures in living Escherichia coli cells by using green fluorescent protein. Proc Natl Acad Sci U S A 93, 12998–13003, doi:10.1073/pnas.93.23.12998 (1996).

5 Hussain, S. et al. MreB filaments align along greatest principal membrane curvature to orient cell wall synthesis. Elife 7, doi:10.7554/eLife.32471 (2018).

6 Kim, S. et al. Microtubule- and dynein-mediated movement of Orientia tsutsugamushi to the microtubule organizing center. Infect Immun 69, 494–500, doi:10.1128/IAI.69.1.494-500.2001 (2001).

7 Keller, C. et al. Dissemination of Orientia tsutsugamushi and inflammatory responses in a murine model of scrub typhus. PLoS Negl Trop Dis 8, e3064, doi:10.1371/journal.pntd.0003064 (2014).

8 Paris, D. et al. Orientia tsutsugamushi in human scrub typhus eschars shows tropism for dendritic cells and monocytes rather than endothelium. PLoS Negl Trop Dis 6, e1466, doi:10.1371/journal.pntd.0001466 (2012).

9 Atwal, S. et al. The obligate intracellular bacterium Orientia tsutsugamushi differentiates into a developmentally distinct extracellular state. Nat Commun 13, 3603, doi:10.1038/s41467-022-31176-9 (2022).

10 Adcox, H. E., Berk, J. M., Hochstrasser, M. & Carlyon, J. A. Orientia tsutsugamushi OtDUB Is Expressed and Interacts with Adaptor Protein Complexes during Infection. Infect Immun, e0046922, doi:10.1128/iai.00469-22 (2022).

11 Luce-Fedrow, A. et al. A Review of Scrub Typhus (Orientia tsutsugamushi and Related Organisms): Then, Now, and Tomorrow. Tropical Medicine and Infectious Disease 3, 8, doi:10.3390/tropicalmed3010008 (2018).

12 Bonell, A., Lubell, Y., Newton, P. N., Crump, J. A. & Paris, D. H. Estimating the burden of scrub typhus: A systematic review. PLoS Negl Trop Dis 11, e0005838, doi:10.1371/journal.pntd.0005838 (2017).

13 Moron, C., Popov, V., Feng, H., Wear, D. & Walker, D. Identification of the target cells of Orientia tsutsugamushi in human cases of scrub typhus. Mod Pathol 14, 752–759, doi:10.1038/modpathol.3880385 (2001).

14 Salje, J. Cells within cells: Rickettsiales and the obligate intracellular bacterial lifestyle. Nat Rev Microbiol, doi:10.1038/s41579-020-00507-2 (2021).

15 Chu, H. et al. Exploitation of the endocytic pathway by Orientia tsutsugamushi in nonprofessional phagocytes. Infect Immun 74, 4246–4253, doi:10.1128/IAI.01620-05 (2006).

16 Schermelleh, L. et al. Subdiffraction multicolor imaging of the nuclear periphery with 3D structured illumination microscopy. Science 320, 1332–1336, doi:10.1126/science.1156947 (2008).

17 Wu, Y. & Shroff, H. Faster, sharper, and deeper: structured illumination microscopy for biological imaging. Nat Methods 15, 1011–1019, doi:10.1038/s41592-018-0211-z (2018).

18 Eggeling, C., Willig, K. I., Sahl, S. J. & Hell, S. W. Lens-based fluorescence nanoscopy. Q Rev Biophys 48, 178–243, doi:10.1017/S0033583514000146 (2015).

19 Vangindertael, J. et al. An introduction to optical super-resolution microscopy for the adventurous biologist. Methods Appl Fluoresc 6, 022003, doi:10.1088/2050-6120/aaae0c (2018).

20 Prakash, K., Diederich, B., Heintzmann, R. & Schermelleh, L. Super-resolution microscopy: a brief history and new avenues. Philos Trans A Math Phys Eng Sci 380, 20210110, doi:10.1098/rsta.2021.0110 (2022).

21 Hell, S. W. & Wichmann, J. Breaking the diffraction resolution limit by stimulated emission: stimulated-emission-depletion fluorescence microscopy. Opt Lett 19, 780–782, doi:10.1364/ol.19.000780 (1994).

22 Klar, T. A. & Hell, S. W. Subdiffraction resolution in far-field fluorescence microscopy. Opt Lett 24, 954–956, doi:10.1364/ol.24.000954 (1999).

23 Klar, T. A., Jakobs, S., Dyba, M., Egner, A. & Hell, S. W. Fluorescence microscopy with diffraction resolution barrier broken by stimulated emission. Proc Natl Acad Sci U S A 97, 8206–8210, doi:10.1073/pnas.97.15.8206 (2000).

24 Huang, B., Bates, M. & Zhuang, X. Super-resolution fluorescence microscopy. Annu Rev Biochem 78, 993–1016, doi:10.1146/annurev.biochem.77.061906.092014 (2009).

25 Heintzmann, R. & Huser, T. Super-Resolution Structured Illumination Microscopy. Chem Rev 117, 13890–13908, doi:10.1021/acs.chemrev.7b00218 (2017).

26 Gustafsson, M. G. et al. Three-dimensional resolution doubling in wide-field fluorescence microscopy by structured illumination. Biophys J 94, 4957–4970, doi:10.1529/biophysj.107.120345 (2008).

27 Kner, P., Chhun, B. B., Griffis, E. R., Winoto, L. & Gustafsson, M. G. Super-resolution video microscopy of live cells by structured illumination. Nat Methods 6, 339–342, doi:10.1038/nmeth.1324 (2009).

28 Muller, C. B. & Enderlein, J. Image scanning microscopy. Phys Rev Lett 104, 198101, doi:10.1103/PhysRevLett.104.198101 (2010).

29 York, A. G. et al. Resolution doubling in live, multicellular organisms via multifocal structured illumination microscopy. Nat Methods 9, 749–754, doi:10.1038/nmeth.2025 (2012).

30 Sheppard, C. Super-resolution in confocal imaging. Optik - International Journal for Light and Electron Optics 80 (1988).

31 Wu, X. & Hammer, J. A. ZEISS Airyscan: Optimizing Usage for Fast, Gentle, Super-Resolution Imaging. Methods Mol Biol 2304, 111–130, doi:10.1007/978-1-0716-1402-0_5 (2021).

32 Rehman, J., Grimm, F., Schloetel, J.-G. & Wurm, C. A. See you at the molecular scale. microscopy Today 29, 34–41 (2021).

33 Lambert, T. J. & Waters, J. C. Navigating challenges in the application of superresolution microscopy. J Cell Biol 216, 53–63, doi:10.1083/jcb.201610011 (2017).

34 Schermelleh, L. et al. Super-resolution microscopy demystified. Nat Cell Biol 21, 72–84, doi:10.1038/s41556-018-0251-8 (2019).

35 Prakash, K. et al. Resolution in super-resolution microscopy - definition, trade-offs and perspectives. Nat Rev Mol Cell Biol, doi:10.1038/s41580-024-00755-7 (2024).

36 Jonkman, J., Brown, C. M., Wright, G. D., Anderson, K. I. & North, A. J. Tutorial: guidance for quantitative confocal microscopy. Nat Protoc 15, 1585–1611, doi:10.1038/s41596-020-0313-9 (2020).

37 Demmerle, J. et al. Strategic and practical guidelines for successful structured illumination microscopy. Nat Protoc 12, 988–1010, doi:10.1038/nprot.2017.019 (2017).

38 Gould, T., Pellett, P. & Bewersdorf, J. in Fluorescence Microscopy: from principles to biological applications (ed U Kubitscheck) Ch. 10s, (WILEY, 2013).

39 Wurm, C. A., Neumann, D., Schmidt, R., Egner, A. & Jakobs, S. Sample preparation for STED microscopy. Methods Mol Biol 591, 185–199, doi:10.1007/978-1-60761-404-3_11 (2010).

40 Buckers, J., Wildanger, D., Vicidomini, G., Kastrup, L. & Hell, S. W. Simultaneous multi-lifetime multi-color STED imaging for colocalization analyses. Opt Express 19, 3130–3143, doi:10.1364/OE.19.003130 (2011).

41 Krishnan, S. & Klingauf, J. The readily retrievable pool of synaptic vesicles. Biol Chem 404, 385–397, doi:10.1515/hsz-2022-0298 (2023).

42 Wurm, C. A. et al. Novel red fluorophores with superior performance in STED microscopy. Optical Nanoscopy 1, 7, doi:10.1186/2192-2853-1-7 (2012).

43 Finzel, L. & Reuss, M. A Stimulated Emission Depletion (STED) Microscope of All Trades. Microscopy Today 30, 26–33 (2022).

44 Spahn, C., Grimm, J. B., Lavis, L. D., Lampe, M. & Heilemann, M. Whole-Cell, 3D, and Multicolor STED Imaging with Exchangeable Fluorophores. Nano Lett 19, 500–505, doi:10.1021/acs.nanolett.8b04385 (2019).

45 Prakash, K. Laser-free super-resolution microscopy. Philos Trans A Math Phys Eng Sci 379, 20200144, doi:10.1098/rsta.2020.0144 (2021).

46 Han, Y. et al. Three-dimensional multi-color optical nanoscopy at sub-10-nm resolution based on small-molecule organic probes. Cell Rep Methods 3, 100556, doi:10.1016/j.crmeth.2023.100556 (2023).

47 Barentine, A. E. S., Schroeder, L. K., Graff, M., Baddeley, D. & Bewersdorf, J. Simultaneously Measuring Image Features and Resolution in Live-Cell STED Images. Biophys J 115, 951–956, doi:10.1016/j.bpj.2018.07.028 (2018).

48 Raab, M. et al. Using DNA origami nanorulers as traceable distance measurement standards and nanoscopic benchmark structures. Sci Rep 8, 1780, doi:10.1038/s41598-018-19905-x (2018).

49 Royon, A. & Converset, N. Quality Control of Fluorescence Imaging Systems. Optik and Photonik 12, 22–25 (2017).

50 Wegel, E. et al. Imaging cellular structures in super-resolution with SIM, STED and Localisation Microscopy: A practical comparison. Sci Rep 6, 27290, doi:10.1038/srep27290 (2016).

51 York, A. G. et al. Instant super-resolution imaging in live cells and embryos via analog image processing. Nat Methods 10, 1122–1126, doi:10.1038/nmeth.2687 (2013).

52 Lee, S. et al. Identification of Outer Membrane Vesicles Derived from Orientia tsutsugamushi. J Korean Med Sci 30, 866–870, doi:10.3346/jkms.2015.30.7.866 (2015).

53 Sliusarenko, O., Heinritz, J., Emonet, T. & Jacobs-Wagner, C. High-throughput, subpixel precision analysis of bacterial morphogenesis and intracellular spatio-temporal dynamics. Mol Microbiol 80, 612–627, doi:10.1111/j.1365-2958.2011.07579.x (2011).

54 Lucas, A. M. et al. Open-source deep-learning software for bioimage segmentation. Mol Biol Cell 32, 823–829, doi:10.1091/mbc.E20-10-0660 (2021).

55 Spahn, C. et al. DeepBacs for multi-task bacterial image analysis using open-source deep learning approaches. Commun Biol 5, 688, doi:10.1038/s42003-022-03634-z (2022).

56 Stringer, C., Wang, T., Michaelos, M. & Pachitariu, M. Cellpose: a generalist algorithm for cellular segmentation. Nat Methods 18, 100–106, doi:10.1038/s41592-020-01018-x (2021).

57 Kleinberg, G., Wang, S., Comellas, E., Monaghan, J. R. & Shefelbine, S. J. Usability of deep learning pipelines for 3D nuclei identification with Stardist and Cellpose. Cells Dev 172, 203806, doi:10.1016/j.cdev.2022.203806 (2022).

58 Atwal, S., Giengkam, S., VanNieuwenhze, M. & Salje, J. Live imaging of the genetically intractable obligate intracellular bacteria Orientia tsutsugamushi using a panel of fluorescent dyes. J Microbiol Methods 130, 169–176, doi:10.1016/j.mimet.2016.08.022 (2016).

59 Mika-Gospodorz, B. et al. Dual RNA-seq of Orientia tsutsugamushi informs on host-pathogen interactions for this neglected intracellular human pathogen. Nat Commun 11, 3363, doi:10.1038/s41467-020-17094-8 (2020).

60 Jahr, W., Velicky, P. & Danzl, J. G. Strategies to maximize performance in STimulated Emission Depletion (STED) nanoscopy of biological specimens. Methods 174, 27–41, doi:10.1016/j.ymeth.2019.07.019 (2020).

61 Boehm, U. et al. QUAREP-LiMi: a community endeavor to advance quality assessment and reproducibility in light microscopy. Nat Methods 18, 1423–1426, doi:10.1038/s41592-021-01162-y (2021).

62 Li, X. et al. Three-dimensional structured illumination microscopy with enhanced axial resolution. Nat Biotechnol 41, 1307–1319, doi:10.1038/s41587-022-01651-1 (2023).

63 Schmidt, U., Weigert, M., Braoaddus, C. & Myers, G. Cell Detection with Star-convex Polygons. arXiv (2018).

64 Wolny, A. et al. Accurate and versatile 3D segmentation of plant tissues at cellular resolution. Elife 9, doi:10.7554/eLife.57613 (2020).

65 Giengkam, S. et al. Orientia tsutsugamushi: comprehensive analysis of the mobilome of a highly fragmented and repetitive genome reveals the capacity for ongoing lateral gene transfer in an obligate intracellular bacterium. mSphere 8, e0026823, doi:10.1128/msphere.00268-23 (2023).

66 Batty EM, C. S., Blacksell SB, Richards A, Paris D, Bowden R, Chan C, Lachumanan R, Day N, Donnelly P, Chen SL, Salje J. Long-read whole genome sequencing and comparative analysis of six strains of the human pathogen Orientia tsutsugamushi. Plos Negl Trop Dis (2018).

67 Giengkam, S. et al. Improved Quantification, Propagation, Purification and Storage of the Obligate Intracellular Human Pathogen Orientia tsutsugamushi. PLoS Negl Trop Dis 9, e0004009, doi:10.1371/journal.pntd.0004009 (2015).

68 Schindelin, J. et al. Fiji: an open-source platform for biological-image analysis. Nat Methods 9, 676–682, doi:10.1038/nmeth.2019 (2012).

69 Brocher, J. biovoxxel/bv3dbox: BioVoxxel 3D Box - v1.22.0 (bv3dbox-1.22.0). Zenodo, doi: 10.5281/zenodo.10982438 (2024).

